# Correlative superresolution microscopy with deep UV reactivation

**DOI:** 10.1101/2023.07.16.549188

**Authors:** Kirti Prakash

## Abstract

Correlative superresolution microscopy has the potential to accurately visualize and validate new biological structures past the diffraction limit. However, combining different superresolution modalities, such as deterministic stimulated emission depletion (STED) and stochastic single-molecule localization microscopy (SMLM), is a challenging endeavor. For correlative STED and SMLM, the following poses a significant challenge: (1) the photobleaching of the fluorophores in STED; (2) the subsequent reactivation of the fluorophores for SMLM; and (3) finding the right fluorochrome and imaging buffer for both imaging modalities. Here, we highlight how the deep ultraviolet (DBUE) wavelengths of the Mercury (Hg) arc lamp can help recover STED bleaching and allow for the reactivation of single molecules for SMLM imaging. We also show that Alexa Fluor 594 and the commercially available Prolong Diamond turn out to be excellent fluorophores and imaging media for correlative STED and SMLM.

## Introduction

The continuous advancement of light microscopy technologies has led to significant milestones in spatial resolution, including confocal microscopy (Minsky, 1988), 4Pi microscopy (Hell et al., 1994), two-photon microscopy (Denk et al., 1990), STED microscopy (Hell and Wichmann, 1994; Baer, 1999), structured illumination microscopy (SIM) (Heintzmann and Cremer, 1999; Gustafsson, 2000), and SMLM (Betzig et al., 2006; Hess et al., 2006; Rust et al., 2006). More recently, correlative light microscopy techniques like MINFLUX, SIMFLUX and RESI have achieved spatial resolutions down to a few nanometers (Balzarotti et al., 2017; Cnossen et al., 2020; Jouchet et al., 2019; Reymond et al., 2020; Reinhardt et al., 2023).

While the allure of achieving resolutions around 1 nm is compelling, the reality is that many biological applications do not require such extreme resolution (Prakash, 2022; Prakash and Curd, 2023; Gwosch et al., 2023). Moreover, fluorophores at distances <10 nm cannot be reliably resolved with SMLM-type methods (Helmerich et al., 2022). Often new biological insights occur in the range of 20-200 nm, highlighting the need for sequential correlative superresolution microscopy. Such correlative imaging using different microscopes with compatible resolution and identical imaging conditions will enable independent validation of findings.

### The experimental setup for laser-free superresolution microscopy (LFSM)

The initial motivation to develop LFSM was to test if single molecules could be detected with an incoherent light source such as a Hg arc lamp and, in turn, reconstruct a superresolution image beyond the diffraction limit (Abbe, 1873; Dickson et al., 1997; Lidke et al., 2005). Traditional super-resolution microscopy techniques employ lasers as the primary light source due to their monochromatic and coherent nature (Betzig et al., 2006). The choice of laser wavelength is optimized for the peak emission of the fluorophore but for single-molecule studies, this is not an essential requirement as long enough photons are extracted for the localization of a fluorophore (Prakash, 2021). Moreover, by successfully implementing the LFSM technique on a conventional microscope, we provide a cost-effective and user-friendly solution for generating high-density, super-resolution images. Our aim is to make super-resolution imaging more accessible to a broader scientific community, even those with limited budgets or access to sophisticated laser-based systems (Hohlbein et al., 2022).

LFSM employs an ordinary epifluorescence microscope (such as Olympus BX61) as found in most cell biology labs. The microscope should ideally have a high NA objective lens (1.4, oil), a CCD camera, and a light source with a minimum power density of 1*kW/cm*^2^. A high-power Hg arc lamp (200 W) would provide the relevant power density (Figure 1A-B). The current LFSM setup uses Alexa 594 as its peak excitation intensity coincides with the excitation wavelengths of the Hg lamp (Figure 1C). Also, due to its high photon count and low duty cycle, Alexa 594 is an excellent fluorophore for performing correlative SMLM and STED measurements (Figure 1I). The deep blue excitation wavelengths (350-380 nm) of the Hg lamp are used to reactivate the blinking of fluorophores when it has slowed down or stopped (Figure 1C). This reactivation allows for continuous imaging and data acquisition (Figure 1G). Commercially available ProLong Diamond is used as the anti-fading mounting media as it works for both STED and LFSM/SMLM imaging. The sample preparation for mouse and amphibian meiotic chromosomes is described in detail here Susiarjo et al. (2009); Prakash et al. (2015). Single-molecule data acquisition, localization, and image reconstruction can be performed with any publicly available SMLM data reconstruction software such as ThunderSTORM (Ovesný et al., 2014).

**Fig. 1.**
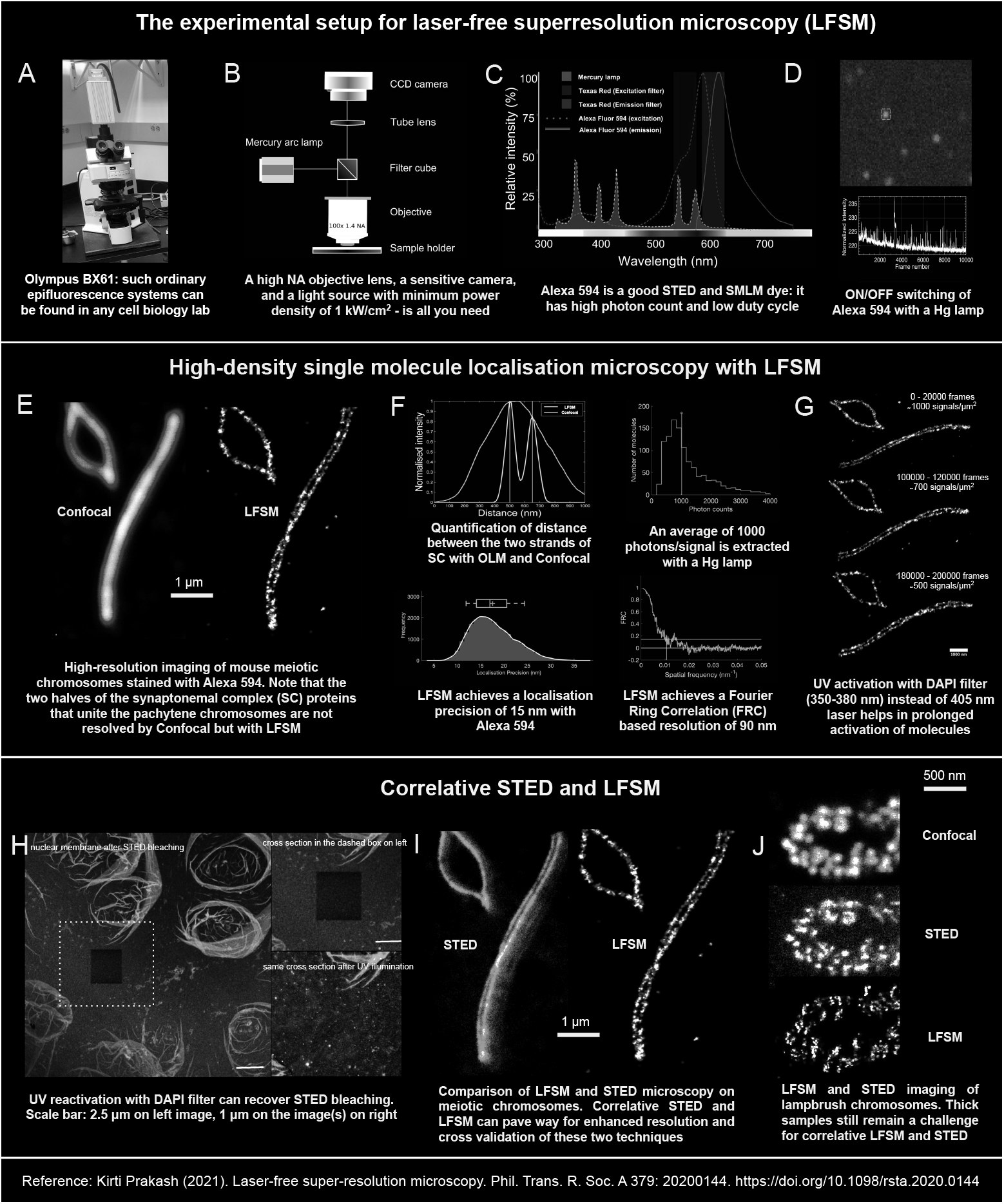
Laser-free superresolution microscopy. (A-D) The experimental setup. (E-G) Achieving high-density localization of single-molecules with LFSM. (H-J) The framework for correlative superresolution microscopy.

### A framework for correlative superresolution microscopy

Based on the LFSM setup, we propose a framework for sequential confocal, STED, and SMLM on the same sample using the same set of fluorophores (Figure 2). This approach allows researchers to obtain complementary information and validate observations across different superresolution techniques, ultimately enhancing our understanding of biological processes.

**Fig. 2.**
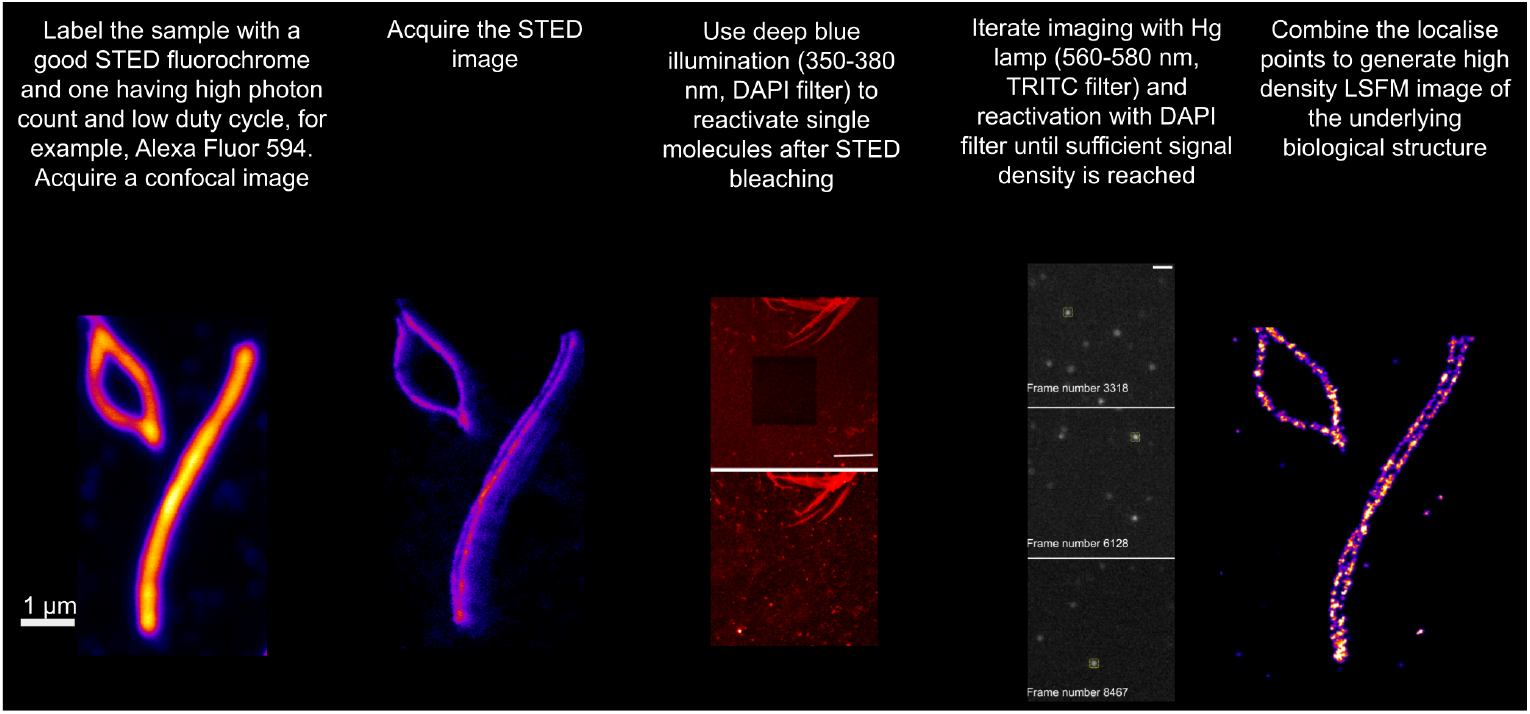
A framework for correlative superresolution microscopy with STED and SMLM/LFSM. Mouse pachytene chromosomes stained with an antibody targeting the synaptonemal complex (SC) are first imaged with a confocal microscope. The two halves of the SC that connect the meiotic chromosome pair remain unresolved by confocal microscope. The same samples are then imaged with a STED and LFSM microscope. The two halves of the SC can now be resolved. While the overall structure (the two strands of SC) remains the same, LFSM reveals more substructure at the nanoscale, highlighting the need of validating new biological findings using methods with similar resolution ranges.

The key to correlative STED and SMLM is the reversal of STED bleaching by UV reactivation using a DAPI excitation filter (365/30 nm) (Figure 1H). The bleached cross-section of the nuclear envelope was successfully recovered following STED acquisition by leveraging the spectral peaks emitted by the Hg arc lamp in the range of 350-380 nm (Figure 1C). Notably, Alexa Fluor 594, known for its brightness and photostability, proved to be suitable for both STED and LFSM imaging. First, we acquired confocal images of Alexa Fluor 594 using a Leica SP8 microscope, followed by STED imaging (Figure 2). Next, we image the same sample region using an Olympus BX 61 microscope in widefield mode. The correlative method helps to identify structural discrepancies at the nanoscale (Figure 1J).

### Conclusion and outlook

The findings presented here highlight the potential of deep blue UV excitation (DBUE) in recovering STED photobleaching, enabling extended imaging and improved sample preservation. LFSM demonstrates the feasibility of ON/OFF switching and localization of single molecules using a standard, unmodified epifluorescence microscope commonly available in cell biology laboratories. Furthermore, the use of a short burst of DBUE (350-380 nm) proves effective in reactivating blinking and allows for the reconstruction of super-resolved images with high signal density. Notably, the combination of Alexa Fluor 594 and a simple imaging buffer, such as Prolong Diamond, enables the sequential imaging of STED and SMLM measurements on the same section of a biological sample.

Our study also demonstrates that high-resolution single-molecule imaging can be achieved using a conventional microscope and incoherent light sources, challenging the notion that lasers and coherent light are indispensable for the on/off switching of single molecules. By utilizing a simple Hg lamp as a non-coherent light source, we provide a cost-effective solution for generating high-density super-resolution images, making this technique accessible to labs with limited budget.

These findings contribute to expanding the repertoire of techniques available for high-density correlative superresolution imaging and pave the way for further advancements in the field. Moreover, the photophysical observations obtained in this study pave the way for more comprehensive investigations into the fundamental processes of fluorophore excitation, photobleaching, and photoactivation.

### Practical considersations

- Use of deep blue UV excitation (DBUE) to recover STED photobleaching
- The ON/OFF switching and localization of single molecules with laser-free superresolution microscopy (LFSM)
- High-density single-molecule imaging using a standard, unmodified epifluorescence microscope commonly found in cell biology laboratories.
- A short burst of DBUE (350–380 nm) to reactivate blinking, once blinking has slowed or stopped. Prolongated blinking helped to reconstruct super-resolved images of high signal density.
- Using Alexa Fluor 594 and a simple imaging buffer (Prolong Diamond) to perform STED and LFSM measurements on the same section of a biological sample.
- The method currently works with flat cell lines and thin samples like chromosome spreads.

### Data, code, and talk availability

- The data and code are available via Zenodo: https://zenodo.org/record/4395254.
- YouTube link to the RMS Imaging ONEWORLDtalk: https://www.youtube.com/watch?v=rfftwIaiUew.
- The primary reference for this article is https://royalsocietypublishing.org/doi/10.1098/rsta.2020.0144.

### Competing Interests

The authors declare that they have no known competing financial interests or personal relationships that could influence the work reported in this paper.

## Biography

Kirti Prakash is a scientist at the Institute of Cancer Research in London, UK. He specializes in the field of super-resolution microscopy and has a research focus on the structure and function of chromosomes.

## Notes

### Competing Interest Statement

The authors have declared no competing interest.

